# CD40 ligation-induced ERK activation promotes autophagy which leads to enhanced radiosensitivity in cervical carcinoma cells

**DOI:** 10.1101/2024.11.18.624076

**Authors:** Baocai Liu, Yadong Zhang, Quan Wang, Qian Wang, Zhixin Wang, Li Feng

**Affiliations:** Department of Radiation Oncology, China-Japan Union Hospital of Jilin University, Changchun, 130033 China; Department of Urology, China-Japan Union Hospital of Jilin University, Changchun, 130033 China

**Keywords:** CD40, autophagy, radiosensitivity, cervical cancer

## Abstract

CD40, a member of the tumor necrosis factor (TNF) receptor superfamily, plays an important role not only in the immune system, but also in tumor progression. CD40 ligation reportedly promotes autophagy in immune cells. However, the effects of CD40 ligation on autophagy and its mechanism in solid tumor cells are still unclear. In this study, we find that CD40 ligation promotes autophagosome formation, and consequently promotes autophagic flux in cervical cancer cells. Mechanically, this effect relies on that ERK contributes to CD40 ligation-induced ATG13 upregulation by p53. Furthermore, we demonstrates that CD40 ligation-induced autophagy increases the radiosensitivity of cervical cancer cells. Taken together, our results provide new evidence for involvement of the CD40 pathway in autophagy and radiotherapy in cervical cancer cells.

## Introduction

CD40 is not only expressed on normal B lymphocytes and antigen presenting cells, but also be expressed on the surface of epithelioid tumors (cervical cancer, ovarian cancer, lung cancer, bladder cancer, liver cancer) and hematological tumors [1, 2]. CD40L (CD154) as a type II membrane protein is a ligand of CD40, which can be divided into two types: soluble and membrane-bound, both of which can exert their biological effects after binding to CD40 [3]. CD40 can form homotrimer under the stimulation of CD40L, or combine with other members of TNFR family to form heterotrimer. Polymerized CD40 can be autophosphorylated to recruit downstream signal molecules, which activate PI3K-AKT, Ras-Raf-MEK-ERK and STAT3 signal pathways to regulate gene expression, thus playing an important role in humoral and cell-mediated immune responses, and participating in the development of tumor. Studies have shown that CD40 activation can inhibit the survival of many kinds of tumor cells, such as B-cell lymphoma, multiple myeloma, bladder cancer, ovarian cancer, breast cancer, skin cancer and cervical cancer, and improve the sensitivity of tumor cells to drugs [4, 5], but its effect on cancer radiosensitivity is still unclear.

Cervical cancer is the fourth most common cancer among women worldwide, with approximately 604000 new cases and 342000 deaths in 2020, of which 90% are in developing countries [6]. At present, about 80% of invasive cervical cancer requires radiotherapy, among which radiotherapy is the main treatment method for IB2-IIA non-surgical patients and IIB-IV patients [7]. Radiotherapy can induce autophagy which can regulate the radiosensitivity. However, the role of autophagy in radiotherapy for cervical cancer is controversial [8, 9]. Some reports indicate that upregulation of autophagy has a cellular protective effect on cervical cancer cells. On the contrary, other studies have shown that induction of autophagy can increase the radiosensitivity of cervical cancer, and the reason for this contradiction is not clear, which may be related to the level of autophagy.

Autophagy flux determines the level of autophagy, which consists of the following successive steps: formation of the omegasome from the endoplasmic reticulum, expansion and nucleation of the phagophore into an autophagosome, maturation into amphisomes by fusing autophagosomes with endosomal compartments, and fusion with lysosomes to form autolysosomes [10–13]. In mammals, the initiation of autophagy is controlled by the autophagy-related protein (ATG)1/ULK1 kinase complex, which contains the Ser/Thr kinase, ULK1, and the accessory proteins ATG13, FIP200, and ATG101 [14, 15]. ATG13 is critical for correctly localizing ULK1 to the pre-autophagosome, and stability of the ULK1 protein [16]. ULK1 kinase activity is essential for the subsequent recruitment of the ATG14-containing class III phosphatidylinositol 3-kinase complex, which leads to the enrichment of phosphatidylinositol 3-phosphate (PtdIns3P) in the endoplasmic reticulum and eventually to the recruitment of PtdIns3P-binding proteins, such as zinc finger, FYVE domain containing 1 (ZFYVE1) to the site of autophagosome generation [17–19]. Two ubiquitin-like conjugation systems, ATG12 and ATG8/LC3B, are required for expansion and completion of autophagosomes [20, 21].

CD40 ligation reportedly promotes autophagy to kill parasites in immune cells [22, 23]. However, The role and mechanism of CD40 in autophagy and radiosensitivity of cervical cancer cells are still unclear. In this study, we demonstrated that CD40 activation enhanced the formation of autophagosomes and autophagic flux by increasing extracellular signal-regulated kinase (ERK)-p53 signaling-mediated upregulation of ATG13 in cervical carcinoma cells, moreover, the CD40 signaling pathway enhanced the radiosensitivity of cervical cancer cells by increasing autophagy level. This provides a new clue for involvement of the CD40 pathway in autophagy and radiotherapy in solid cancer cells.

## Materials and Methods

### Cell culture and transfections

Human HeLa and SiHa cells were purchased from the American Type Culture Collection (Manassas, VA, USA) and were grown, respectively, in RPMI-1640 and Dulbecco’s modified Eagle’s medium supplemented with 10% fetal bovine serum. The pcDNA3/CD40 plasmid was transfected into HeLa cells using Lipofectamine 2000 (Thermo Fisher Scientific, Waltham, MA, USA). The cells were cultured in medium containing 500 µg/mL G418 for two weeks at 37°C for selection. HeLa cells stably overexpressing CD40 (HeLa/CD40 cells) were treated with 500 ng/mL CD40 ligand (CD40L/CD154) (Cell Signaling Technology, Beverly, MA, USA) for 24 h or pretreated for 30 min with 10 µM U0126 (Cell Signaling Technology) before being stimulated with CD40L. The stably transfected GFP-LC3B HeLa cell line (HeLa/GFP-LC3B cells) was a kind gift from Yingyu Chen (Peking University, Beijing, China). Autophagy was inhibited by treating cells with 25 μM chloroquine (Sigma-Aldrich, St. Louis, MO, USA) for 4 h. Cells were transfected with plasmids using MegaTran 1.0 Transfection Reagent (OriGene, Rockville, MD, USA) and with small interfering RNAs (siRNAs) using Lipofectamine 2000.

### Plasmid construction and siRNA

The GFP-ZFYVE1 and mTagRFP-mWasabi-LC3B plasmids were kindly provided by Yingyu Chen. The mCherry-ATG5 and CD40 plasmids were constructed by our laboratory. All plasmids were confirmed by DNA sequencing. siRNAs targeting CD40 were designed and synthesized by Qiagen (Germantown, MD, USA), and siRNAs targeting ATG13 were from GenePharma (Suzhou, China). The siRNA sequences for CD40 and ATG13 are listed in Table S1.

### Semiquantitative real-time polymerase chain reaction (RT-PCR)

Total RNA samples from control and CD154-stimulated cells were extracted with TRIzol reagent (Invitrogen, Grand Island, NY, USA). RT-PCR was performed using the ThermoScript RT-PCR System (Invitrogen). The primers for key autophagy genes are listed in Table S2.

### Western blot analyses

Co-immunoprecipitation assays were performed with control and CD40L-stimulated HeLa/CD40 cells. After 24 h of CD40L stimulation, cells were harvested in a buffer containing 20 mM Tris-HCl (pH 8.0), 150 mM NaCl, 2 mM EDTA, 10% glycerol, 0.5% Nonidet P-40, 1 mM dithiothreitol, 1 mM phenylmethylsulfonyl fluoride, 5 μg/mL leupeptin, 5 μg/mL aprotinin, 5 μg/mL pepstatin, and 1% protease inhibitor cocktail (Roche, Basel, Switzerland). Protein concentrations were determined using the bicinchoninic acid assay (Pierce, Rockford, IL, USA). Whole cell lysates were fractionated using 10% sodium dodecyl sulfate polyacrylamide gel electrophoresis and electrotransferred onto polyvinylidene difluoride membranes (GE Healthcare). Western blotting was performed according to standard protocols [24].

The following antibodies were used: anti-GFP, anti-GAPDH, anti-ATG13, anti-ULK1, anti-p62, anti-ERK, anti-phospho-ERK^Thr202/Tyr204^ (all from Cell Signaling Technology), anti-LC3B (Sigma-Aldrich), and anti-CD40 (Abcam, Cambridge, MA, USA). Secondary antibodies included anti-mouse and anti-rabbit DyLight 800- and DyLight 680-conjugated IgG (Rockland Antibodies & Assays, Limerick, PA, USA). Signals were detected by an Odyssey Infrared Imager (LI-COR Bioscience, Lincoln, NE, USA) or viewed using ECL Plus (GE Healthcare).

### Confocal microscopy

Cells were washed with phosphate-buffered saline, fixed, and permeabilized with 4% paraformaldehyde for 30 min at 4°C. They were then blocked in a solution containing 0.1% Triton X-100 and incubated with the indicated antibodies for 60 min at 37°C. The cells were then washed twice with phosphate-buffered saline and stained with Hoechst 33342 (Cell Signaling Technology) for 10 min before being imaged with an Ultra View VOX confocal laser scanning microscope (Perkin Elmer, Waltham, MA, USA).

### Colony formation experiment

Cells at logarithmic growth phase were seeded into a six well plate with a certain number of cells (300 cells/well in the 0Gy group, 1500 cells/well in the 2Gy group, 3000 cells/well in the 4Gy group, 6000 cells/well in the 6Gy group, and 8000 cells/well in the 8Gy group), with three replicates in each group. According to the experimental requirements, cells were irradiated with different doses and cultured for 2 weeks. The conditioned medium (containing 500ng/ml CD40L) was changed every 3 days to observe the formation of clones. Cells were fixed and stained with 0.5% crystal violet (Beyotime Institute of Biotechnology), and colonies of at least 50 cells were counted by GelCount (Oxford Optronix, Oxfordshire, Great Britain). Adhesion rate=number of clones per well in the control group/number of cells inoculated per well, Survival fraction (SF)=number of clones per well in the experimental group/(number of cells implanted per well x adhesion rate)*100%. Survival curves were fitted in a multi-target single-hit model [SC=C1C−C(1C−Ce−D/D0)N] using Graphpad Prism 5.0 (Graphpad Software, San Diego, CA, USA).

### Statistical analyses

Data are expressed as means ± SEM. Statistical analyses were performed using two-tailed Student’s *t*-tests in Prism 5.0 (Graphpad Software, San Diego, CA, USA). Differences were considered significant if *P* < 0.05.

## Results

### CD40 ligation promotes autophagic flux

To evaluate whether CD40 ligation induced autophagy in cervical cancer cells, we first analyzed the expression of endogenous LC3B-II, a marker of autophagosomes, and the level of p62, an autophagy cargo receptor, in HeLa/CD40 cells by western blot. CD40 ligation increased the conversion of LC3B-I to LC3B-II and decreased p62 expression (Fig. 1A).

**Fig. 1.**
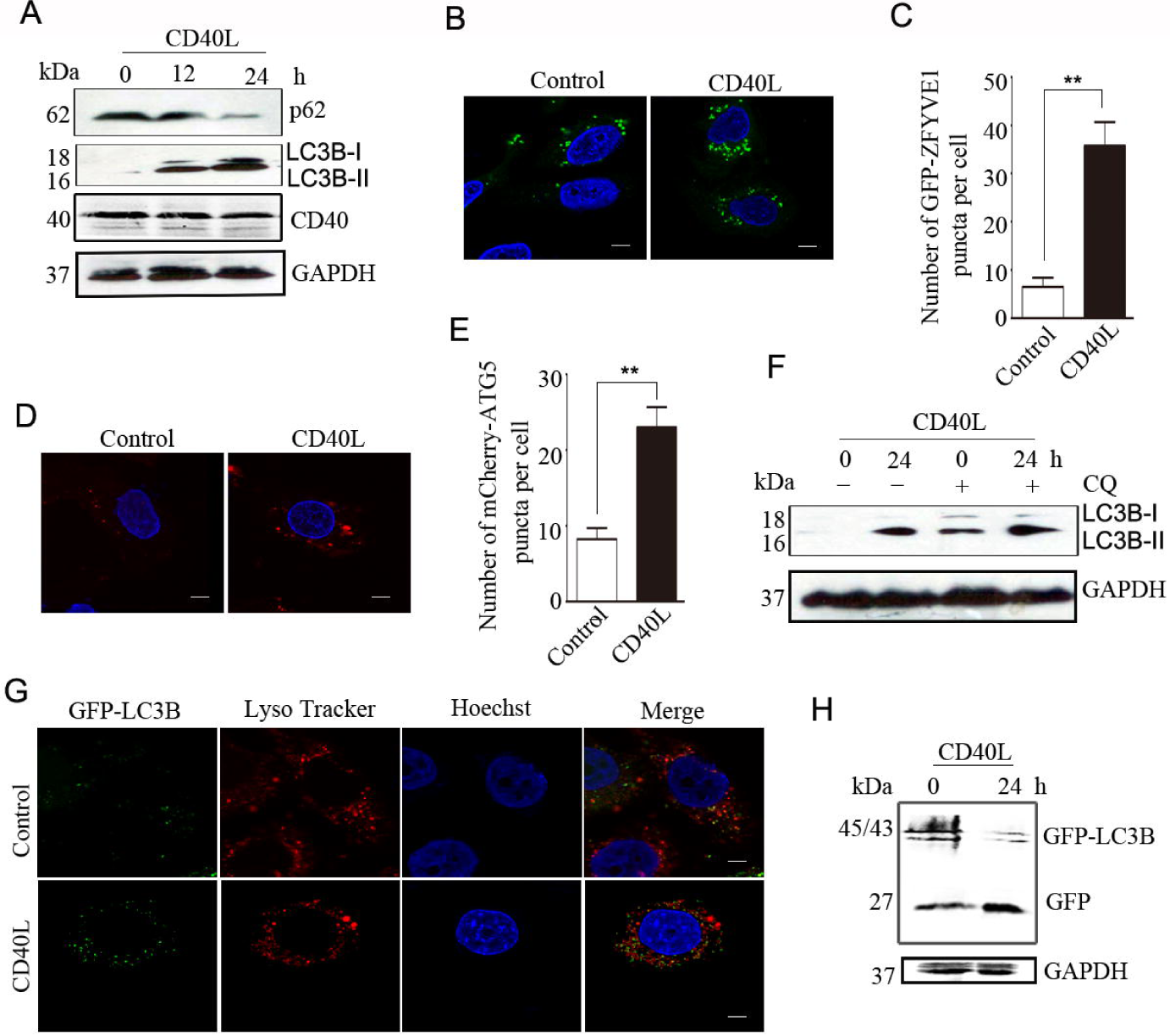
CD40 ligation promotes autophagic flux. **(A)** HeLa/CD40 cells were treated with CD40L for the indicated times. The levels of LC3B, p62, and CD40 proteins were detected by western blot. GAPDH was used as the internal standard. **(B)** HeLa/CD40 cells were transfected with GFP-ZFYVE1 for 12 h, then stimulated with CD40L for 24 h. After fixation, cell images were captured using a confocal microscope. Scale bars: 5 µm. **(C)** GFP-ZFYVE1 puncta per cell were quantified. The results are expressed as means ± SEM of at least 20 cells (\*\**P* < 0.01). **(D)** HeLa/CD40 cells were transfected with mCherry-ATG5, then treated as in (A). Cell images were captured using a confocal microscope. Scale bars: 5 µm. (**E)** mCherry-ATG5 puncta per cell were quantified. The results are expressed as means ± SEM of at least 20 cells (\*\**P* < 0.01). **(F)** HeLa/CD40 cells were stimulated with CD40L for 24 h, then treated with 25 µM chloroquine (CQ) for 4 h. The level of LC3B protein was detected by western blot. GAPDH was used as the internal standard. **(G)** HeLa/GFP-LC3B cells were transfected with a CD40 plasmid. Representative fluorescence microscopy images of the co-localization of GFP-LC3B with LysoTracker Red 24 h after transfection are shown. Scale bars: 5 µm. (**H)** HeLa/GFP-LC3B cells were transfected with a CD40 plasmid. The levels of free GFP were analyzed by western blot. Data are representative of three independent experiments.

To determine the step at which the autophagy process was affected by CD40 ligation in HeLa/CD40 cells, we examined the formation of omegasomes that can be marked specifically by the endoplasmic reticulum-associated PtdIns3P-binding protein, ZFYVE1. HeLa/CD40 cells transfected to overexpress GFP-ZFYVE1 were observed under a confocal microscope. CD40 ligation significantly increased the number of GFP-ZFYVE1-labeled vesicles (Fig. 1B, C), suggesting that CD40 ligation enhanced the formation of omegasomes.

Subsequently, we detected the formation of ATG5-labeled membrane structures, which are autophagosome precursors. As shown in Figure 1D and E, CD40 ligation increased the number of membrane-bound mCherry-ATG5 structures in HeLa/CD40 cells. These results indicate that CD40 ligation may enhance autophagosome formation.

To confirm these results, the lysosome inhibitor, chloroquine, was used to prevent fusion between autophagosomes and lysosomes. In the presence of chloroquine, more LC3B-II was accumulated in CD40L-stimulated than control cells (Fig. 1F). This indicated that the elevated LC3B-II levels driven by CD40 ligation were from increased autophagosome formation.

We then overexpressed CD40 in HeLa/GFP-LC3B cells and used LysoTracker Red to label lysosomes. More GFP-LC3B puncta were observed to co-localize with lysosomes in CD40L-stimulated than control HeLa/GFP-LC3B cells (Fig. 1G), suggesting that CD40 activation may facilitate autophagic flux. After delivery to lysosomes, the GFP-LC3B protein will be cleaved and LC3B is rapidly degraded, while the GFP moiety remains stable [25]. In HeLa/GFP-LC3B cells, the free GFP level was upregulated by CD40L stimulation compared to control as illustrated by western blot (Fig. 1H). Taken together, these data support the hypothesis that CD40 activation promotes autophagic flux by increasing autophagosome formation.

### Knockdown of CD40 impairs CD40L-mediated autophagy

To confirm the effects of CD40 ligation on autophagy regulation, we used CD40 siRNAs to knock down endogenous CD40 in SiHa cells. CD40 levels were significantly silenced by siCD40-1, 2, and 3, as shown in Figure 2A and B. CD40 ligation-stimulated LC3B-II expression was examined by western blot and found to be decreased in CD40 knockdown compared with control cells (Fig. 2C). Consistent with this, the p62 level was increased in CD40L-treated CD40 knockdown cells (Fig. 2C).

**Fig. 2.**
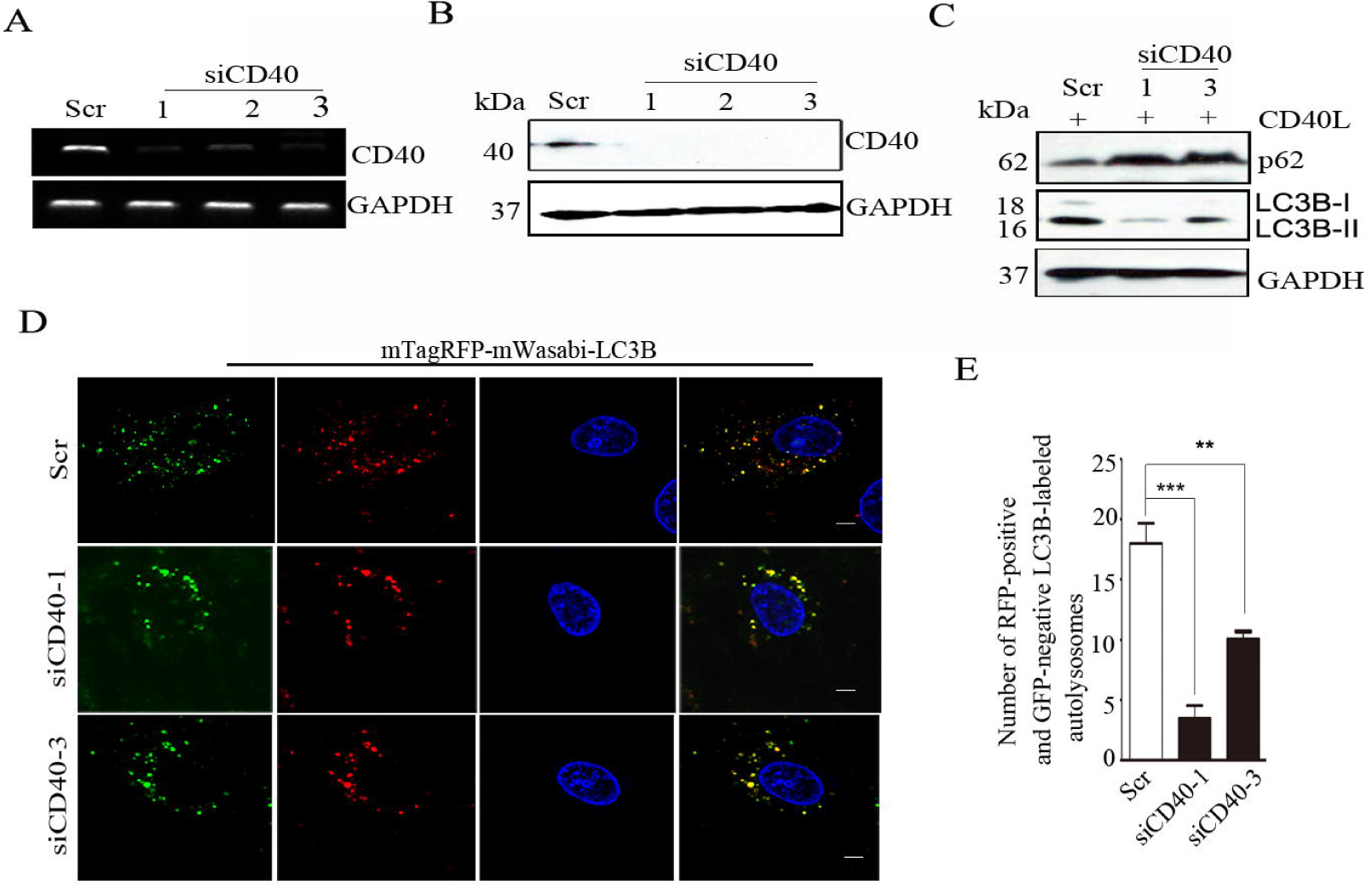
Knockdown of CD40 impairs CD40L-mediated autophagy. SiHa cells were transfected with control siRNA (Scr), or siCD40-1, 2, or 3. CD40 knockdown was detected by the semi-quantitative polymerase chain reaction (**A)** and western blot (**B)**. (**C**) SiHa cells were transfected with Scr, or siCD40-1 or -3, and stimulated with CD40L for 24 h. The expression of LC3B and p62 proteins was analyzed by western blot. GAPDH was used as the internal standard. **(D)** Representative images of mTagRFP-mWasabi-LC3B distribution in CD40L-stimulated SiHa cells co-transfected with mTagRFP-mWasabi-LC3B and Scr or siCD40-1. Scale bar: 5 µm. (**E**) The number of RFP-positive and GFP-negative LC3B-labeled autolysosomes per cell was quantified. The results are expressed as means ± SEM of at least 20 cells (\*\**P* < 0.01, \*\*\**P* < 0.001).

The mTagRFP-mWasabi-LC3B reporter was used to evaluate autophagic flux in CD40 knockdown SiHa cells. As shown in Figure 2D and E, CD40 knockdown decreased the CD40L-induced transition of mTagRFP-mWasabi-LC3B-positive autophagosomes to mTagRFP-positive, mWasabi-negative autolysosomes. Taken together, these results suggest that CD40 activation facilitates autophagy in cervical carcinoma cells.

### CD40 ligation promotes autophagy by upregulating ATG13 expression

The core ATG proteins involved in autophagosome formation are divided into five subgroups: ATG1/ULK1 protein-kinase complex, ATG9-ATG2-ATG18 complex, Vps34-ATG6/Beclin 1 class III phosphatidylinositol 3-kinase complex, and the ATG12 and ATG8/LC3 conjugation systems. Because CD40 activation can affect gene transcription through its downstream signaling pathways, we first detected the mRNA expression of core ATG genes in CD40L-stimulated HeLa/CD40 cells. The results showed that, among the ATG molecules examined, only the transcription of ATG13 was increased by CD40 ligation (Fig. 3A, B). Western blot analysis demonstrated that CD40 ligation also increased expression of the ATG13 protein (Fig. 3C, D). The effect of CD40 ligation on ATG13 expression was confirmed in CD40 knockdown SiHa cells (Fig. 3E, F). These results indicate that CD40 ligation increases ATG13 expression.

**Fig. 3.**
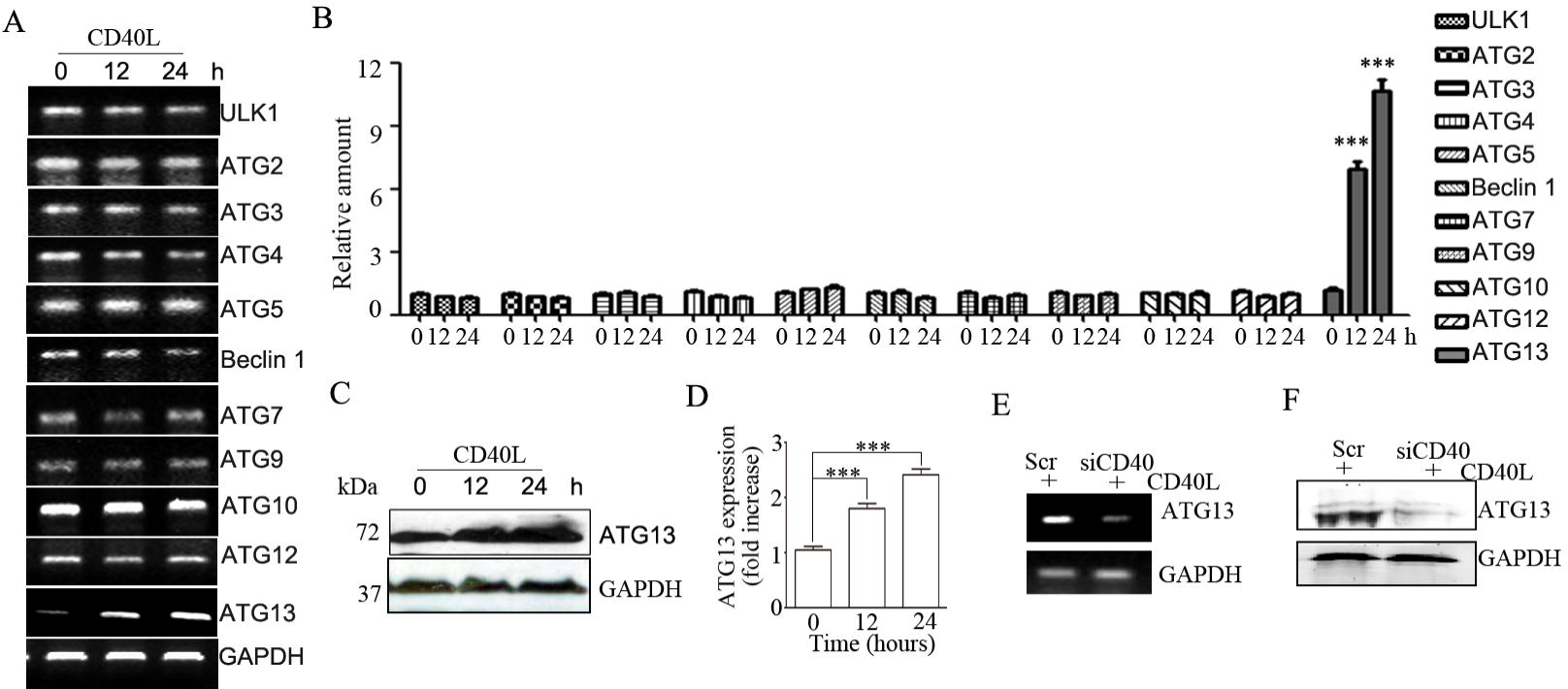
CD40 ligation increases ATG13 expression. **(A)** HeLa/CD40 cells were treated with CD40L for the indicated times. The mRNA expression of core ATG genes was analyzed using the semi-quantitative polymerase chain reaction. (**B)** The grey densities of the target bands were analyzed by ImageJ software and normalized to the grey density of GAPDH. The average relative grey density with SEM from three independent experiments was shown (\*\*\**P*<0.001). (**C)** The protein expression of ATG13 was analyzed by western blot. **(D)** The grey densities of the target bands were analyzed by ImageJ software and normalized to the grey density of GAPDH. The average relative grey density with SEM from three independent experiments was shown (\*\*\**P*<0.001). SiHa cells were transfected with control siRNA (Scr), or siCD40-1, and stimulated with CD40L for 12h. The expression of ATG13 was analyzed by the semi-quantitative polymerase chain reaction (**E)** and western blot (**F)**. GAPDH was used as the internal standard.

To test whether CD40 ligation-stimulated autophagy was dependent on ATG13 expression, we used several ATG13 siRNAs to knock down endogenous ATG13 in HeLa/CD40 cells, and assessed LC3B-II expression, showing that ATG13 levels were significantly silenced by siATG13-1 and 3 (Fig. 4A, B). Compared with control siRNA cells, knockdown of ATG13 decreased CD40 ligation-stimulated LC3B-II expression (Fig. 4C, D). In CD40 overexpressed HeLa/GFP-LC3B cells, knockdown of ATG13 reversed the CD40 ligation-stimulated formation of GFP-LC3B puncta (Fig. 4E, F) and the upregulated free GFP level by CD40L stimulation (Fig. 4G). These results suggest that CD40 ligation promotes autophagy by increasing ATG13 expression.

**Fig. 4.**
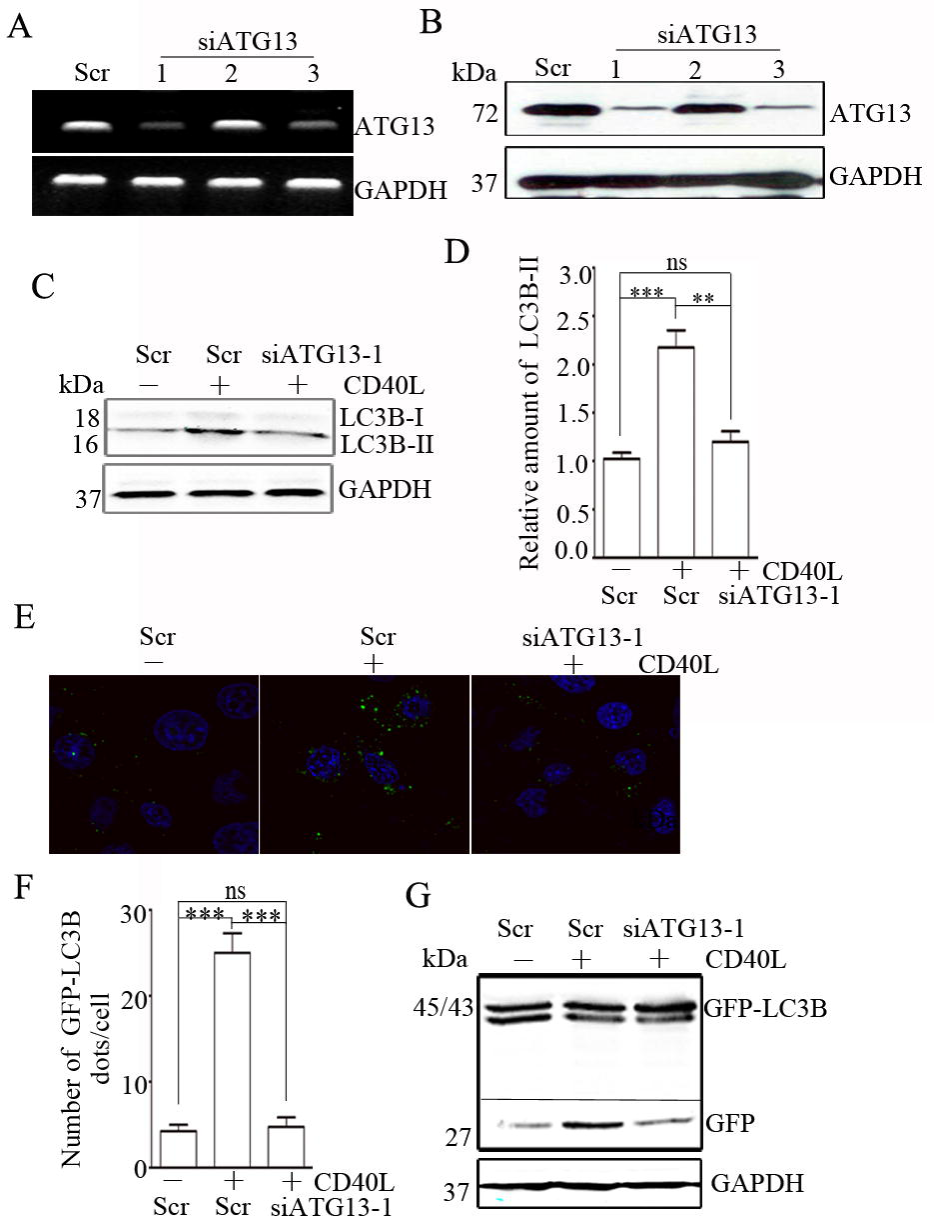
ATG13 mediates CD40 ligation-induced autophagy. HeLa/CD40 cells were transfected with three distinct ATG13 shRNAs or the matching control non-targeting siRNA (Scr). ATG13 levels were detected by the semi-quantitative polymerase chain reaction (**A)** and western blot (**B)**. (**C)** HeLa/CD40 cells were transfected with Scr or siATG13-1. The expression of LC3B protein was analyzed by western blot. GAPDH was used as the internal standard. (**D)** The grey densities of the target bands were analyzed by ImageJ software and normalized to the grey density of GAPDH. The average relative grey density with SEM from three independent experiments was shown (\*\*\**P*<0.001). (**E)** Representative confocal microscopy images of GFP-LC3B distribution were shown. (**F)** GFP-LC3B puncta per cell were quantified and expressed as the mean ± SEM of at least 20 cells (\*\*\**P*<0.001, ns, not significant). (**G)** The levels of free GFP were analyzed by western blot.

### CD40 ligation increases ERK-p53 signal transduction

CD40 ligation induces the mitogen-activated protein kinase signaling pathways to affect the expression of a large number of genes [26]. We found that CD40 ligation increased the phosphorylation level of ERK in HeLa/CD40 cells, but not p38 (Fig. 5A, sFig. 1). Importantly, the ERK inhibitor, U0126, abrogated the upregulated expression of ATG13 and LC3B-II induced by CD40 ligation (Fig. 5B, C). This suggested that ERK signaling contributed to CD40 ligation-induced autophagy and ATG13 upregulation.

**Fig. 5.**
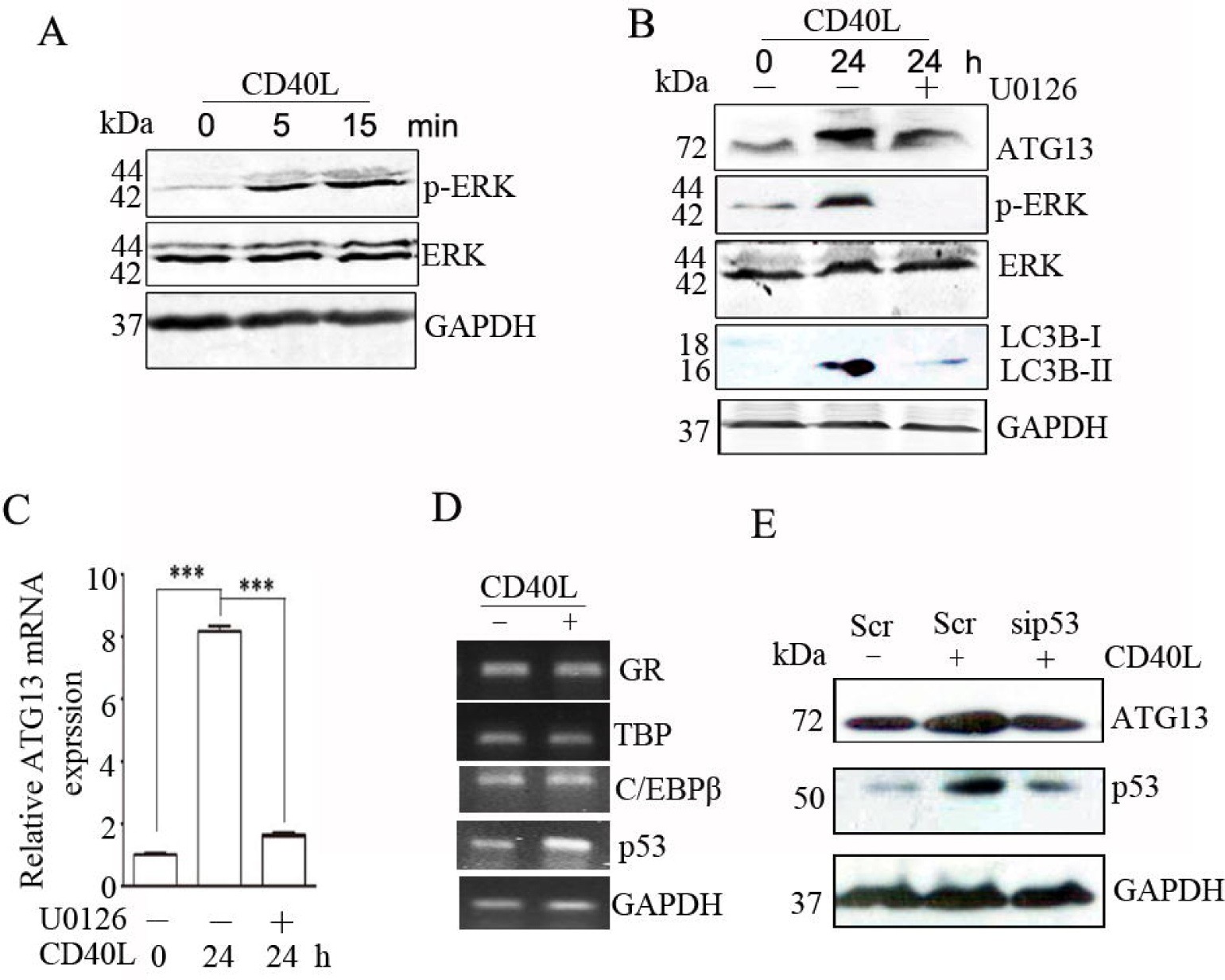
CD40 ligation increases ERK-p53 signal transduction. **(A)** HeLa/CD40 cells were stimulated with CD40L for the indicated times. The levels of phosphorylated and total ERK were detected by western blot. (**B)** HeLa/CD40 cells were treated with CD40L for 24 h or pretreated with U0126 for 30 min before being stimulated with CD40L. The levels of ATG13, LC3B, phosphorylated ERK, and total ERK were detected by western blot, and ATG13 mRNA expression was analyzed using real-time RT-PCR(**C**). (**D)** HeLa/CD40 cells were stimulated with CD40L for 12 h, C/EBPβ, p53, TBP and GR mRNA levels were detected by the semi-quantitative polymerase chain reaction. (**E)** HeLa/CD40 cells were transfected with Scr or sip53, after 24 h, cells were treated with CD40L for 12 h, the levels of ATG13 and p53 were detected by western blot. Data are representative of three independent experiments.

In order to determine the transcription factor that CD40-ERK regulates the expression of ATG13, we first predicted the transcription factors in the functional region of the ATG13 promoter by using the UCSC database (http://genome.ucsc.edu/) and the PROMO database (http://alggen.lsi.upc.es/). A total of 24 transcription factors were found (data not shown). By reviewing the literature, we further selected the transcription factors regulated by ERK/MAPK signal pathway, including C/EBPβ, p53, TBP and GR [27–31], for further study. We found that CD40 ligation increased the transcription level of p53, but not C/EBP β, TBP and GR in HeLa/CD40 cells (Fig. 5D), which was confirmed in CD40 knockdown SiHa cells (sFig. 2). Futhermore, consistent with the transcription level, CD40 ligation also increased p53 expression in protein level (Fig. 5E). Importantly, p53 knockdown reversed the enhanced effect of CD40 ligation on ATG13 expression (Fig. 5E). The above results indicated that p53 mediates an increase in ATG13 levels in CD40-ERK signal transduction.

### CD40 ligation-induced autophagy increases radiosensitivity of cervical cancer cells

We have demonstrated that CD40 activation can increase the autophagy level of cervical cancer cells. We further detected the effect of CD40 ligation on radiation-induced autophagy through western blot. As shown in Figure 6A and B, CD40 ligation can further increase the level of radiation-induced LC3B-II, indicating that CD40 activation can promote radiation-induced autophagy. Recent studies have shown that high expression of CD40/CD40L is associated with a better prognosis of cervical cancer [32], but whether CD40 activation affects the radiosensitivity of cervical cancer cells is unclear. The survival curve of a multi-target single-hit model simulated from clone formation experiments showed that CD40 activation increased the radiosensitivity of cervical cancer HeLa/CD40 cells, which can be reversed by autophagy inhibitor CQ (Fig. 6C), and the same effect was obtained in SiHa cells (sFig. 3), suggesting that CD40/CD40L-induced autophagy enhances radiosensitivity of cervical cancer cells.

**Fig. 6.**
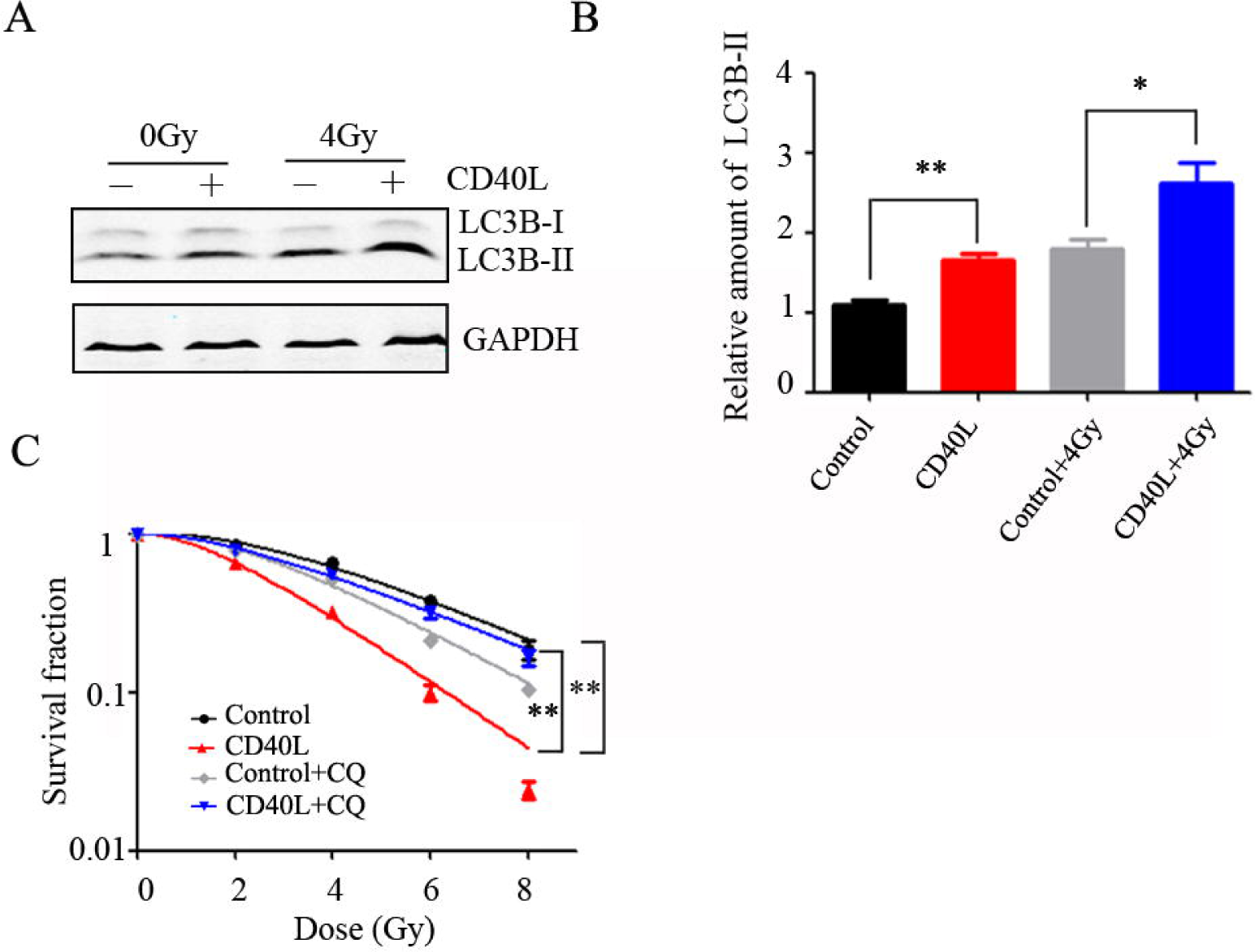
CD40 ligation-induced autophagy increases radiosensitivity of cervical cancer cells. **(A)** HeLa/CD40 cells were treated with CD40L or 4Gy X-ray alone or combination for 24 h. The levels of LC3B and GAPDH proteins were detected by western blot. Normalized densitometry data using ImageJ software according to the mean values of three independent experiments was presented as meansC±Cs.d. (\**P*C<C0.05, \*\**P*C<C0.01) **(B)**. **(C)** Clonogenic survival fraction curves of control, CD40L, control+CQ and CD40L+CQ groups following exposure to 0, 2, 4, 6, 8CGy of X-rays (\*\**P*C<C0.01).

## Discussion

Accumulating evidence suggests that CD40 activation can increase autophagy through different molecular mechanisms to kill *Toxoplasma gondii.* For example, in macrophages, CD40 activation enhanced autophagic flux to kill *T. gondii* through increasing ULK1 phosphorylation and dissociating Bcl-2 and Beclin 1 [22, 23]. However, the functional effect of CD40 activation on autophagy in solid tumor cells was still unclear.

In this paper, we used cervical carcinoma cells and demonstrated that CD40 ligation enhanced the formation of autophagosome, consequently promoting autophagic flux by increasing ATG13 expression. In our experimental system, CD40 ligation increased only the mRNA level of ATG13 among the eleven examined ATG molecules. ATG13 can function as an adaptor through recruiting ULK1, FIP200, and ATG101 to form the ULK1 protein kinase complex. Furthermore, ATG13 is critical for stabilizing the ULK1 protein in 293T, HeLa and mouse embryonic fibroblast cells [33–35]. In the current study, we found that CD40 ligation had no effect on the mRNA level of ULK1 in HeLa cells, but increased its protein level (sFig. 4). These results suggested that CD40 activation may increase the stability of ULK1 by regulating ATG13 expression.

As a tumor suppressor, p53 coordinates a variety of responses mainly by regulating the transcription of its target genes, including cell cycle arrest, DNA repair, aging and apoptosis [36, 37]. Studies have also proved that p53 can induce autophagy directly as a transcription factor of autophagy-related genes. For example, p53 upregulates the transcription of Cathepsin D, TGM2, Sestrin1, Sestrin2, BNIP3, and further upregulates autophagy [36]; Combined ChIP sequencing and RNA sequencing analysis revealed that p53 binds to many autophagy genes, including ATG2, ATG4, ATG7 and ATG10 [38], but whether p53 affects the autophagy gene ATG13 has not been reported. In this study, we found that p53, as the downstream of CD40-ERK signal pathway, directly upregulated the transcriptional expression of ATG13 to increase autophagy, suggesting that p53 is a transcription factor of ATG13, which needs further experimental verification.

The effect of autophagy on the radiosensitivity of cervical cancer has been reported, but the results are inconsistent. For example, radiotherapy can increase the autophagy level of HTB35 cervical cancer cells, and inhibiting autophagy by knocking down autophagy-related genes (ATG3, ATG12) can increase the sensitivity of the cells to radiotherapy, indicating that radiotherapy-induced autophagy can reduce radiosensitivity [8]. However, recent research has shown that the radiosensitivity of cervical cancer can be elevated by photobio modulation (PBM) through autophagy pathways which lead to the induction of apoptosis, increased ROS and damaged DNA [9]. In the current study, activating CD40 signaling enhanced autophagic flux and promoted the radiosensitivity of cervical cancer cells which can be reversed by autophagy inhibitor. It is worth noting that in our experimental system, autophagy induced solely by radiotherapy plays a protective role in cervical cancer cells. In recent years, Dr. Gerwitz’s lab found that dual functions of autophagy in the response of breast tumor cells to radiation: cytoprotective autophagy with radiation alone and cytotoxic autophagy in radiosensitization by vitamin D3 [39]. Based on this, they proposed an attractive theory about the “autophagy switch”, which describes autophagy as a cytoprotective process in irradiated breast tumor cells. However, after increasing autophagy to a certain extent, protective autophagy can transform into cytotoxic autophagy, thereby increasing radiosensitivity [40]. As in our experimental system, both cytoprotective and cytotoxic autophagy can occur simultaneously in cervical cancer cells, and the underlying mechanisms deserve further study.

In summary, our study demonstrated that CD40 activation was associated with ATG13 expression and autophagy by increasing the phosphorylation level of ERK in cervical carcinoma cells. In addition, the promotion of ATG13 expression was induced by p53. Importantly, we demonstrated that CD40 ligation-induced autophagy increased the radiosensitivity of cervical cancer cells. These results revealed a novel mechanism and function for CD40 ligation as a positive regulator of cell autophagy in cervical carcinoma cells, providing new directions for examining the effect of CD40 ligation on autophagy and function in other solid cancer cells.

## Acknowledgements

This work was supported by a grant from the Project of Science and Technology Department of Jilin Province (YDZJ202401242ZYTS)

## References

[1] Yan C, Saleh N, Yang J, Nebhan CA, Vilgelm AE, Reddy EP, Roland JT, Johnson DB, Chen SC, Shattuck-Brandt RL, Ayers GD, Richmond A, Novel induction of CD40 expression by tumor cells with RAS/RAF/PI3K pathway inhibition augments response to checkpoint blockade, Mol Cancer. 20 (2021) 85.

[2] Yan C, Richmond A, Hiding in the dark: pan-cancer characterization of expression and clinical relevance of CD40 to immune checkpoint blockade therapy, Mol Cancer. 20 (2021) 146

[3] Ma DY, Clark EA, The role of CD40 and CD154/CD40L in dendritic cells, Semin Immunol. 21 (2009) 265–72.

[4] Li DK, Wang W, Characteristics and clinical trial results of agonistic anti-CD40 antibodies in the treatment of malignancies, Oncol Lett. 20 (2020) 176.

[5] Bereznaya NM, Chekhun VF, Expression of CD40 and CD40L on tumor cells: the role of their interaction and new approach to immunotherapy, Exp Oncol. 29 (2007) 2–12.

[6] World Health Organization (WHO) Human papillomavirus (HPV) and cervical cancer. Fact sheet. 2022

[7] Faye MD, Alfieri J, Advances in Radiation Oncology for the Treatment of Cervical Cancer, Curr Oncol. 29 (2022) 928–944.

[8] Apel A, Herr I, Schwarz H, Rodemann HP, Mayer A, Blocked autophagy sensitizes resistant carcinoma cells to radiation therapy, Cancer Res. 68 (2008) 1485–94.

[9] Djavid GE, Bigdeli B, Goliaei B, Nikoofar A, Hamblin MR, Photobiomodulation leads to enhanced radiosensitivity through induction of apoptosis and autophagy in human cervical cancer cells, J Biophotonics. 10 (2017) 1732–1742.

[10] Yang Z, Klionsky DJ, Mammalian autophagy: core molecular machinery and signaling regulation, Curr Opin Cell Biol. 22 (2010) 124–31.

[11] Lamark T, Johansen T, Mechanisms of Selective Autophagy, Annu Rev Cell Dev Biol. 37 (2021) 143–169.

[12] Nähse V, Schink KO, Stenmark H, ATPase-regulated autophagosome biogenesis, Autophagy. 20 (2024) 218–219.

[13] Zhen Y, Stenmark H, Autophagosome Biogenesis, Cells. 12 (2023) 668.

[14] Kannangara AR, Poole DM, McEwan CM, Youngs JC, Weerasekara VK, Thornock AM, Lazaro MT, Balasooriya ER, Oh LM, Soderblom EJ, Lee JJ, Simmons DL, Andersen JL, BioID reveals an ATG9A interaction with ATG13-ATG101 in the degradation of p62/SQSTM1-ubiquitin clusters, EMBO Rep. 22 (2021) e51136.

[15] Thorne RF, Yang Y, Wu M, Chen S, TRIMming down autophagy in breast cancer, Autophagy. 18 (2022) 2512–2513.

[16] Alers S, Wesselborg S, Stork B, ATG13: just a companion, or an executor of the autophagic program? Autophagy. 10 (2014) 944–56.

[17] Obara K, Ohsumi Y, Atg14: a key player in orchestrating autophagy, Int J Cell Biol, 2011 (2011) 713435.

[18] Hurley JH, Young LN, Mechanisms of Autophagy Initiation, Annu Rev Biochem, 86 (2017) 225–244.

[19] Chang HC, Tao RN, Tan CT, Wu YJ, Bay BH, Yu VC, The BAX-binding protein MOAP1 associates with LC3 and promotes closure of the phagophore, Autophagy. 17(2021) 3725–3739.

[20] Alam JM, Maruyama T, Noshiro D, Kakuta C, Kotani T, Nakatogawa H, Noda NN, Complete set of the Atg8-E1-E2-E3 conjugation machinery forms an interaction web that mediates membrane shaping, Nat Struct Mol Biol. 31(2024) 170–178.

[21] Klionsky DJ, Abdel-Aziz AK, Abdelfatah S, Abdellatif M, Abdoli A, Abel S, Abeliovich H, et al, Guidelines for the use and interpretation of assays for monitoring autophagy, Autophagy. (2021) 1–382.

[22] Liu E, Lopez Corcino Y, Portillo JA, Miao Y, Subauste CS, Identification of Signaling Pathways by Which CD40 Stimulates Autophagy and Antimicrobial Activity against Toxoplasma gondii in Macrophages, Infect Immun. 84 (2016) 2616–26.

[23] Subauste CS, Andrade RM, Wessendarp M, CD40-TRAF6 and autophagy-dependent anti-microbial activity in macrophages, Autophagy. 3 (2007) 245–8.

[24] Liu B, Su Y, Li T, Yuan W, Mo X, Li H, He Q, Ma D, Han W, CMTM7 knockdown increases tumorigenicity of human non-small cell lung cancer cells and EGFR-AKT signaling by reducing Rab5 activation, Oncotarget. 6 (2015) 41092–107.

[25] Xia D, Qu L, Li G, Hongdu B, Xu C, Lin X, Lou Y, He Q, Ma D, Chen Y, MARCH2 regulates autophagy by promoting CFTR ubiquitination and degradation and PIK3CA-AKT-MTOR signaling, Autophagy. 12 (2016) 1614–30.

[26] Elgueta R, Benson MJ, de Vries VC, Wasiuk A, Guo Y, Noelle RJ, Molecular mechanism and function of CD40/CD40L engagement in the immune system, Immunol Rev. 229 (2009) 152–72.

[27] Zhang T, He YM, Wang JS, Shen J, Xing YY, Xi T, Ursolic acid induces HL60 monocytic differentiation and upregulates C/EBPβ expression by ERK pathway activation, Anticancer Drugs. 22 (2011) 158–65.

[28] Gao M, Zhao LR, Turning Death to Growth: Hematopoietic Growth Factors Promote Neurite Outgrowth through MEK/ERK/p53 Pathway, Mol Neurobiol. 55 (2018) 5913–5925.

[29] Zhong S, Zhang C, Johnson DL, Epidermal growth factor enhances cellular TATA binding protein levels and induces RNA polymerase I- and III-dependent gene activity, Mol Cell Biol. 24 (2004) 5119–29.

[30] Santalucía T, Sánchez-Feutrie M, Felkin LE, Bhavsar PK, Barton PJ, Zorzano A, Yacoub MH, Brand NJ, Phenylephrine requires the TATA box to activate transcription of GLUT1 in neonatal rat cardiac myocytes, J Mol Cell Cardiol. 38 (2005) 677–84.

[31] Numakawa T, Odaka H, Adachi N, Chiba S, Ooshima Y, Matsuno H, Nakajima S, Yoshimura A, Fumimoto K, Hirai Y, Kunugi H, Basic fibroblast growth factor increased glucocorticoid receptors in cortical neurons through MAP kinase pathway, Neurochem Int. 118 (2018) 217–224.

[32] Grazia GA, Bastos DR, Villa LL, CD40/CD40L expression and its prognostic value in cervical cancer, Braz J Med Biol Res. 56 (2023) e13047.

[33] Jung CH, Jun CB, Ro SH, Kim YM, Otto NM, Cao J, Kundu M, Kim DH, ULK-Atg13-FIP200 complexes mediate mTOR signaling to the autophagy machinery, Mol Biol Cell. 20 (2009) 1992–2003.

[34] Ganley IG, Lam du H, Wang J, Ding X, Chen S, Jiang X, ULK1.ATG13.FIP200 complex mediates mTOR signaling and is essential for autophagy, J Biol Chem. 284 (2009) 12297–305.

[35] Hosokawa N, Hara T, Kaizuka T, Kishi C, Takamura A, Miura Y, Iemura S, Natsume T, Takehana K, Yamada N, Guan JL, Oshiro N, Mizushima N, Nutrient-dependent mTORC1 association with the ULK1-Atg13-FIP200 complex required for autophagy, Mol Biol Cell. 20 (2009) 1981–91.

[36] Wang H, Guo M, Wei H, Chen Y, Targeting p53 pathways: mechanisms, structures, and advances in therapy, Signal Transduct Target Ther. 1 (2023) 92.

[37] Xie K, Liu L, Wang M, Li X, Wang B, Yin S, Chen W, Lin Y, Zhu X, IMPA2 blocks cervical cancer cell apoptosis and induces paclitaxel resistance through p53-mediated AIFM2 regulation, Acta Biochim Biophys Sin (Shanghai). 55 (2023) 623–632.

[38] Broz D, Mello S, Bieging K, Jiang D, Dusek R, Brady C, Sidow A, Attardi L, Global genomic profiling reveals an extensive p53-regulated autophagy program contributing to key p53 responses, Genes Dev. 1 (2013) 1016–31.

[39] Wilson EN, Bristol ML, Di X, Maltese WA, Koterba K, Beckman MJ, Gewirtz DA, A switch between cytoprotective and cytotoxic autophagy in the radiosensitization of breast tumor cells by chloroquine and vitamin D, Horm Cancer. 2 (2011) 272–85.

[40] Ondrej M, Cechakova L, Durisova K, Pejchal J, Tichy A, To live or let die: Unclear task of autophagy in the radiosensitization battle, Radiother Oncol. 119 (2016) 265–75.

