## supplementary data for "CD40 ligation-induced ERK activation promotes autophagy which leads to enhanced radiosensitivity in cervical carcinoma cells"

**Supplementary Material**

**Table S1. siRNA sequences.**

| siRNA | Sequence |
| --- | --- |
| Control siRNA | 5′-UUCUCCGAACGUGUCACGUTT-3′ |
| si-CD40-1 | 5′-UUGGCAUCCAUGAAAGUCUC-3′ |
| si-CD40-2 | 5′-UUGGAGAAGAAGCCGACUGGG-3′ |
| si-CD40-3 | 5′-UUUGAUAAAGACCAGCACCAA-3′ |
| si-ATG13-1 | 5′-GGUUCAACUUAGCAAUCAATT-3′ |
| si-ATG13-2 | 5′-GCUGCUGGCUUAAAUGCUATT-3′ |
| si-ATG13-3 | 5′-GCAGCCUCUCCAUAGAUAUTT-3′ |

**Table S2. PCR primer sequences.**

| Primer | Sequence |
| --- | --- |
| ULK1 Forward | 5'-GGCAGTTCTTTGTTCAAGCGTTCC-3' |
| ULK1 Reverse | 5'-CCTCTAACCTGGCCCAGTCCTC-3' |
| ATG2 Forward | 5'-GAGCAGCAGACGGTATTTCGG-3' |
| ATG2 Reverse | 5'-AGACACGCAGGCAAGGGACA-3' |
| ATG3 Forward | 5'-AGTCCACCACTGTCCAACAT-3' |
| ATG3 Reverse | 5'-GTATCTACCCATCCGCCATC-3' |
| ATG4 Forward | 5'-GAGGTGCCCTGCCCATGTTT-3' |
| ATG4 Reverse | 5'-AGCGGGCGGGATGGATGAAA-3' |
| ATG5 Forward | 5'-GCTTCCTGGAGTCCTGCTAC-3' |
| ATG5 Reverse | 5'-TTCCCTTTCAGTTATCTCATCC-3' |
| Beclin 1 Forward | 5'-ACCGCAAGATAGTGGCAGAA-3' |
| Beclin 1 Reverse | 5'-GCGACCCAGCCTGAAGTTAT-3' |
| ATG7 Forward | 5'-GGGGATTTCTTTCACGGTTTG-3' |
| ATG7 Reverse | 5'-TCAGCAGCTTGGGTTTCTTG-3' |
| ATG9 Forward | 5'-TGCTACCTCCGCCACTTCAA-3' |
| ATG9 Reverse | 5'-TGCTCAGGGCAGAACACCAT-3' |
| ATG10 Forward | 5'-AAAGGACTGTTCTGATGGCTAC-3' |
| ATG10 Reverse | 5'-TGCCCAAGTATTGGATGTTC-3' |
| ATG12 Forward | 5'-TGAGGGTTCAGATGATAGAC-3' |
| ATG12 Reverse | 5'-ATTGTGCCATTACAGTCCAG-3' |
| ATG13 Forward | 5'-AGCAATCAAAGACATCCCAGAG-3' |
| ATG13 Reverse | 5'-AAGCCTTCTCCTAAGCCACT-3' |

**sFig. 1 CD40 activation has no effect on p38 phosphorylation.** HeLa/CD40 cells were stimulated with CD40L for the indicated times. The levels of phosphorylated and total p38 were detected by western blot. GAPDH was used as the internal standard.

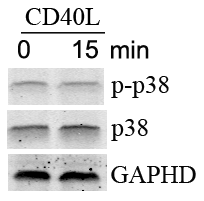

**sFig. 2 Effect of CD40 activation on C/EBPβ, p53, TBP and GR mRNA expression.** SiHa cells were transfected with control siRNA (Scr), or siCD40-1, after 24 h, cells were treated with CD40L for 12 h, C/EBPβ, p53, TBP and GR mRNA levels were detected by the semi-quantitative polymerase chain reaction

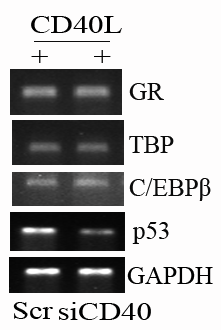

**sFig. 3 CD40 activation increases the radiosensitivity of cervical cancer SiHa cells.** Clonogenic survival fraction curves of control, CD40L, control+CQ and CD40L+CQ groups following exposure to 0, 2, 4, 6, 8 Gy of X-rays in SiHa cells (***P* < 0.01).

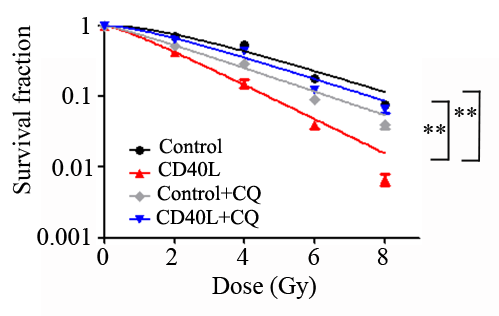

**sFig. 4 CD40 ligation increases ULK1 expression.** HeLa/CD40 cells were treated with CD40L for 24 h. The protein expression of ULK1 was analyzed by western blot.

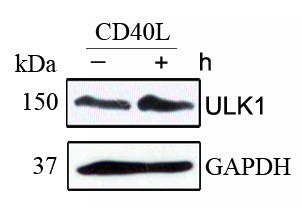
